# Tracking the Fidelity of Internal Neural Representations with Error-In-Variables Regression

**DOI:** 10.64898/2026.04.22.720005

**Authors:** Isabel Garon, Stephen Keeley, Alex H. Williams

## Abstract

Internal neural representations can systematically deviate from externally measured sensory and behavioral variables, yet neuroscientists lack a principled statistical framework to quantify these mismatches. Here we introduce a nonlinear error-in-variables regression framework that explicitly models neural activity as a function of latent internal variables that deviate from measured sensory and behavioral variables. This approach uses a flexible basis expansion and a sampling-based inference scheme to jointly infer neuron-specific tuning functions, latent trajectories, and a representational fidelity parameter *κ* that controls the strength of coupling between latent and measured variables. On synthetic datasets, the model accurately recovers latent dynamics, tuning curves, and identifies the true fidelity regime via cross-validated marginal likelihood. Applied to population recordings from mouse anterodorsal thalamic nucleus and rat medial entorhinal cortex across distinct sensory and behavioral conditions, the framework reveals condition-dependent changes in representational fidelity, tuning gain and profile, and uncovers latent population manifolds that are obscured in conventional tuning analyses. These results establish error-in-variables regression as a powerful and computationally tractable tool for tracking the fidelity of internal neural representations in systems neuroscience experiments.

## 1 Introduction

Neuroscientists are interested in identifying sensory and behavioral variables that correlate with neural activity. However, these correlations are often imperfect constructs. For example, internal representations may systematically deviate from their intended target, such as when navigational circuits accumulate path integration errors over time (Zhang et al. 2014). Moreover, neural populations may represent external variables with a systematic bias or offset, such as when place cells in hippocampus and grid cells in medial entorhinal cortex fire as if representing an animal’s past or future location (Vollan et al. 2025; Frank et al. 2000; Battaglia et al. 2004; Johnson and Redish 2007; Chaudhuri-Vayalambrone et al. 2023; Ouchi and Fujisawa 2024) or spontaneously switch between different spatial reference frames (Kelemen and Fenton 2010). Finally, experimental measurements of sensory inputs and behavioral outputs often contain noise, particularly in naturalistic and unconstrained environments. For example, video tracking of body pose may be error-prone and nominally identical sensory stimuli may elicit distinct responses due to subtle differences in eye, ear, or body position across trials.

For all of these reasons and more, we expect population-level neural representations to systematically diverge from their externally measured counterparts. However, there is no widely agreed upon methodology to measure these discrepancies, even though methods that address this need would be useful in a variety of settings. In working memory tasks, these methods could help diagnose the root cause of error trials by identifying whether discrepancies in representations manifest early in sensory areas or in higher-order cognitive areas (Wimmer et al. 2014; Alleman et al. 2024). In spatial navigation tasks, these methods could provide a moment-by-moment account of coding precision for position, speed, and direction, providing insight into how neural representations support planning and exploration under uncertainty (Chersi and Burgess 2015). In multisensory cue combination tasks, these tools could help assess the quality of neural representations arising from different streams of sensory evidence (Fetsch et al. 2013).

Several existing modeling approaches characterize relationships between neural activity and external variables, but each comes with critical limitations. Supervised sensory encoding models, such as generalized linear models, typically treat externally measured variables as noise-free input variables. While this assumption is attractive from a practical standpoint (e.g., because it enables one to leverage convex optimization methods; Paninski 2004), we will see that it is often violated and therefore can result in misleading or incomplete conclusions. Unsupervised manifold inference models (Wu et al. 2017; Chaudhuri et al. 2019; Jensen et al. 2020; Gardner et al. 2022) have complementary strengths and weaknesses. While they can extract low-dimensional latent variables from neural activity alone, these models are challenging to tune, particularly in data-limited regimes or when neural representations have complex, multi-dimensional geometry (Narayanan and Mitter 2010; Genovese et al. 2012). Furthermore, there is no guarantee that latent variables extracted from a fully unsupervised analysis are easy to interpret.

Here, we develop a framework based on *error-in-variables (EIV) regression* (Carroll et al. 2006) to capture the moment-to-moment mismatch between internal representations and externally measured variables. The fundamental strength of this framework is its ability to continuously interpolate between a fully supervised neural encoding model and a fully unsupervised manifold inference model, allowing practitioners to infer (e.g. through cross-validation) the optimal degree of model supervision for each dataset they consider. Through example analyses of simulated datasets and simultaneously recorded neural populations in the mouse anterodorsal thalamic nucleus (ADn) and rat medial entorhinal cortex (MEC), we argue that EIV regression is a natural and widely applicable model for systems neuroscientists, enabling them to rigorously characterize errors in neural representations on a moment-by-moment basis.

## 2 Results

### 2.1 Error-in-Variables Regression Model

The EIV regression model applies to any experiment where multiple neurons are recorded simultaneously with an external stimulus or behavioral measurement. Formally, let *Y*_*nt*_ denote the activity (e.g. spike count) of neuron *n* at timebin *t*, and let ***s*** _*t*_ denote a vector of sensory or behavioral covariates (e.g. the position and heading direction of the animal). Often, neuroscientists are interested in computing the tuning curve for each neuron, which describes how the expected neural response changes as a function of the stimulus or behavior. The tuning curve is a function *f*_*n*_(·) which is fit to data such that the approximation *Y*_*nt*_ ≈ *f*_*n*_(***s*** _*t*_) holds as well as possible across all neurons and timebins.

The central premise of EIV regression is that external measurements are inexact predictors of neural circuit activity. For example, ***s*** _*t*_ may contain measurement noise—e.g., due to video tracking errors of an animal’s position and body pose. Moreover, neural representations may not match external reality—e.g. an animal’s internal sense of position and heading direction may deviate over time due to path integration errors. To address these sources of error or “noise” in ***s*** _*t*_, we propose to augment traditional tuning curve analysis with an error term ***ϵ***_*t*_ such that neural activity is modeled according to *Y*_*nt*_ ≈ *f*_*n*_(***s*** _*t*_ + ***ϵ***_*t*_).

While intuitive, it is nontrivial to implement EIV regression models. Fitting the model involves simultaneously estimating *f*_*n*_(·) for each neuron and ***ϵ***_*t*_ for each timebin. Classical methods for EIV regression analysis (see Fuller 1987) often make strict assumptions (e.g. linear relationships and gaussian noise) which are violated in neuroscience applications. We therefore developed a nonlinear latent variable model which leverages Gaussian process regression and sampling based approximations of the latent posterior (see *Methods*). In its simplest form, this model outputs an estimate of each neuron’s nonlinear tuning curve, *f*_*n*_(·), as well as the *latent representation* of the neural population at each time bin, denoted ***x*** _*t*_ = ***s*** _*t*_ + ***ϵ***_*t*_ . Because the model is formulated in Bayesian terms, it is possible to go further and obtain approximate posterior distributions over these quantities. However, we focus our narrative on point estimates for simplicity.

To gain intuition on how this model can be used, Figure 1 illustrates three simulated scenarios of a population of neurons encoding head direction. In this case, ***s*** _*t*_ is a one-dimensional circular variable ranging from 0 to 2*π*. In fig. 1a-c, we plot the animal’s latent representation of head direction, ***x*** _*t*_, over time (black line) as well as the measured heading direction, ***s*** _*t*_ (grey dots). In the first scenario (fig. 1a), the latent representation closely matches measured behavior, meaning that ***ϵ***_*t*_ ≈ 0 and therefore ***x*** _*t*_ ≈ ***s*** _*t*_ . The second scenario (fig. 1b) and third scenario (fig. 1c) show simulations with increasing mismatch between ***x*** _*t*_ and ***s*** _*t*_ . The spiking activity, and therefore the latent representation (black line), is exactly preserved across all three simulations to aid visual comparison.

**Figure 1.**
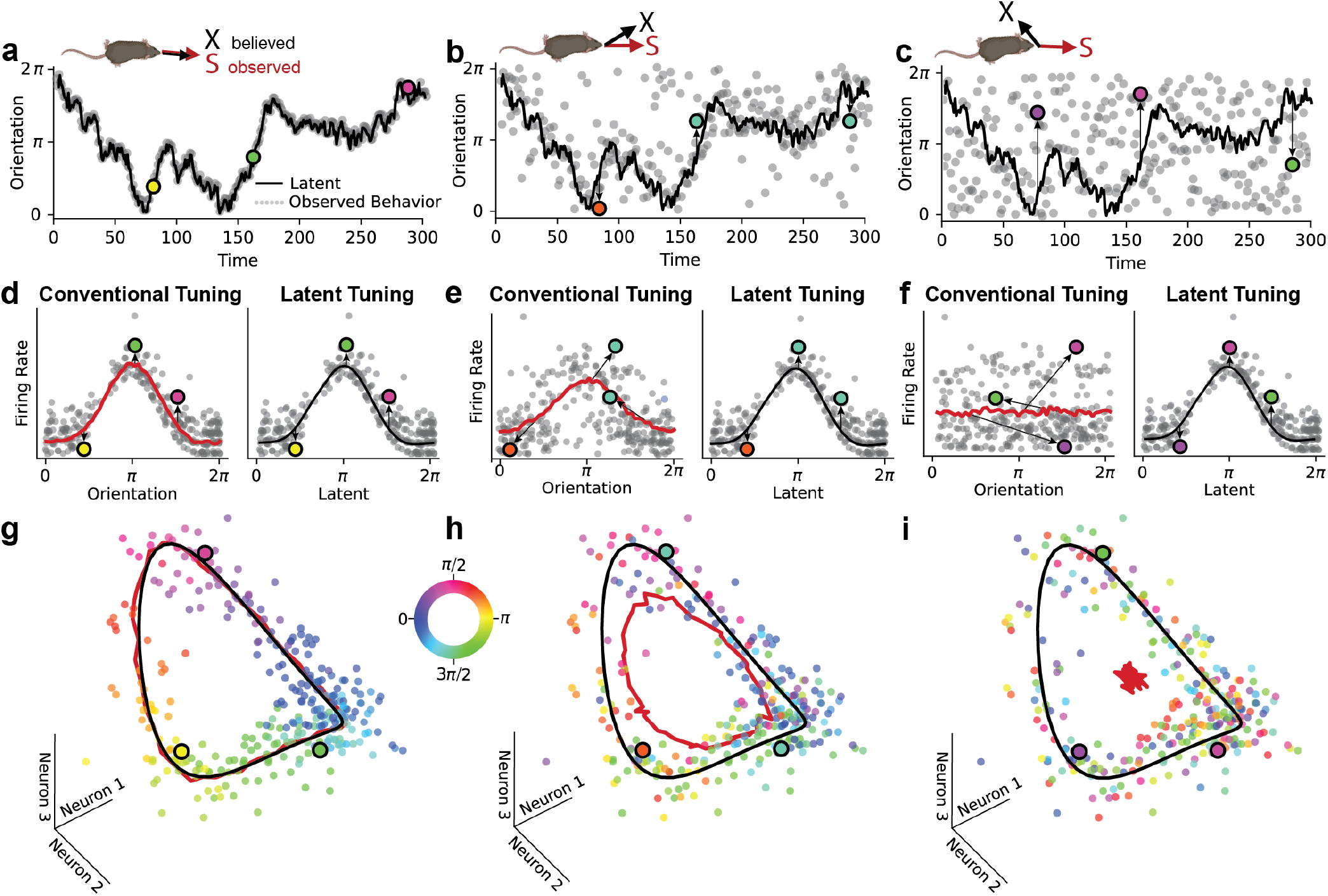
Conceptual Explanation of EIV Regression Model. A-C) Schematic of observed **s** (gray) and latent **x** (black) behavior when **s** and **x** are (A) perfectly aligned, (B) aligned but noisy, and (C) uncorrelated. Latent is identical across all three settings. The behavior (A-C) and firing rates (D-I) for three example time points are highlighted throughout the figure, colored according to observed behavior. D-F) Simulated example of a neuron tuned to the latent variable in A-C. Conventional tuning curves (red) are calculated using the observed behavior. As noise in these observations increases (E, F, arrows), the conventional tuning begins to flatten. Note that the firing rate for each of the example points is identical, it is only position on the x-axis which is changing. Tuning to the latent remains sharp, irrespective of observations. G-I) In each of these three cases, the structure of the neural activity (binned time points represented by scattered dots) remains the same, but the measured behavior varies (color of the dots corresponds to behavioral labels). Tuning to the latent (black) characterizes this population manifold well across all three settings, while conventional tuning (red) based on behavioral labels fails to reflect the true structure of the data, collapsing to the population mean firing rate.

Figure 1d-f show tuning curves of an example neuron across these three scenarios. Grey dots represent measured neural responses, while red lines plot tuning with respect to ***s*** _*t*_ (“conventional tuning”) and black lines plot tuning with respect to ***x*** _*t*_ (“latent tuning”). As described above, the simulated neuron is tuned to ***x*** _*t*_, while ***s*** _*t*_ is an experimentally measurable, but error-prone, proxy for this latent representation. Deviations from the tuning-predicted firing rate along the vertical axis arise solely from neural encoding noise, while deviations along the horizontal axis arise solely from mismatches between ***x*** _*t*_ and ***s*** _*t*_, which we expect to arise for numerous reasons as described in the *Introduction*.

When ***x*** _*t*_ tends to closely match ***s*** _*t*_, we say that the system has high *representational fidelity*. In this case, the conventional tuning curve computed with respect to ***s*** _*t*_ closely matches the ground truth tuning with respect to ***x*** _*t*_ (fig. 1d). When the differences between ***x*** _*t*_ and ***s*** _*t*_ tend to be large, we say that the system has low representational fidelity. In these scenarios, the conventional tuning curves flatten out and poorly predict the neural response (fig. 1e-f). Intuitively, this occurs because the conventional tuning profile does not account for noise along the horizontal axis, while the latent tuning profile, in contrast, is aligned to the appropriate variable (see colored datapoints and arrows indicating deviations from tuning in fig. 1d-f).

The premise of EIV regression is to recover the latent neural tuning curves with respect to ***x*** _*t*_ . Figure 1g-i illustrates why this is possible when multiple, simultaneously recorded neurons are tuned to the same latent representation. Here, 3D scatterplots show simulated activity measurements of three co-recorded neurons. Each point is colored by the measured heading direction, ***s*** _*t*_ . As before, the latent representation and neural responses are identical across the three simulations. Therefore, the 3D position of the datapoints is the same across all three panels.

The joint activity of this simulated neural population lies close to a 1-dimensional ring manifold defined by the latent tuning curves (fig. 1g-i, black curve). When the measured head direction, ***s*** _*t*_, closely matches the latent representation, ***x*** _*t*_, this manifold can be identified by conventional tuning curve analysis (fig. 1g; red curve). However, as the mismatch between ***x*** _*t*_ and ***s*** _*t*_ increases, the manifold identified by conventional tuning curve analysis deviates from the desired latent manifold (fig. 1h). At extreme levels of noise, the conventional tuning curves flatten out and the manifold shrinks to a single point—the mean population firing rate (fig. 1i).

The three simulated scenarios in Figure 1 call for three different statistical modeling strategies. On one extreme, when ***x*** _*t*_ ≈ ***s*** _*t*_, one can simply fit conventional tuning curves. On the other extreme, when ***x*** _*t*_ and ***s*** _*t*_ are strongly mismatched, it would be prudent to ignore ***s*** _*t*_ entirely and attempt to identify the ring structure from the neural data alone. In principle, this can be accomplished by unsupervised manifold learning methods; however, we will see that these models can be challenging to fit when data is limited and when manifolds are complex.

In many cases, neuroscientists are faced with intermediate cases between these two extremes and similar to the scenario outlined in panels B, E, and H of Figure 1. In these situations, one should intuitively leverage the information in ***s*** _*t*_ about ***x*** _*t*_ to construct a better estimate of the latent neural manifold. The EIV regression framework presented here formalizes this intuition, establishing a rigorous method for decoding latent neural manifolds in these varied scenarios.

### 2.2 Demonstration on Synthetic Data

To validate the EIV framework, we first assessed its ability to recover latent neural representations from synthetic datasets characterized by varying degrees of behavioral and sensory noise. Similar to Figure 1, we consider a simple case where ***x*** _*t*_ and ***s*** _*t*_ are 1-dimensional circular variables. The latent variable, ***x*** _*t*_ is noisily encoded by a population of *N* = 30 neurons. See *Methods* for additional simulation details.

Here, we employ an EIV model in which the conditional distribution of ***s*** _*t*_ given ***x*** _*t*_ follows a von Mises distribution centered at ***x*** _*t*_ with concentration *κ*. Thus, *κ*, quantifies the representational fidelity of the system. Systems with high representational fidelity (large *κ*) have low levels of mismatch between ***x*** _*t*_ and ***s*** _*t*_ . Systems with low representational fidelity (small *κ*) have high levels of mismatch.

We simulated data from this generative model at three levels of representational fidelity corresponding to low, medium, and high values of *κ* (fig. 2A; *κ* = 0.5, blue; *κ* = 3.0, yellow; *κ* = 8.0, red). As in Figure 1, we observe that conventional tuning curves degrade as mismatch increases (fig. 2B, black lines).

**Figure 2.**
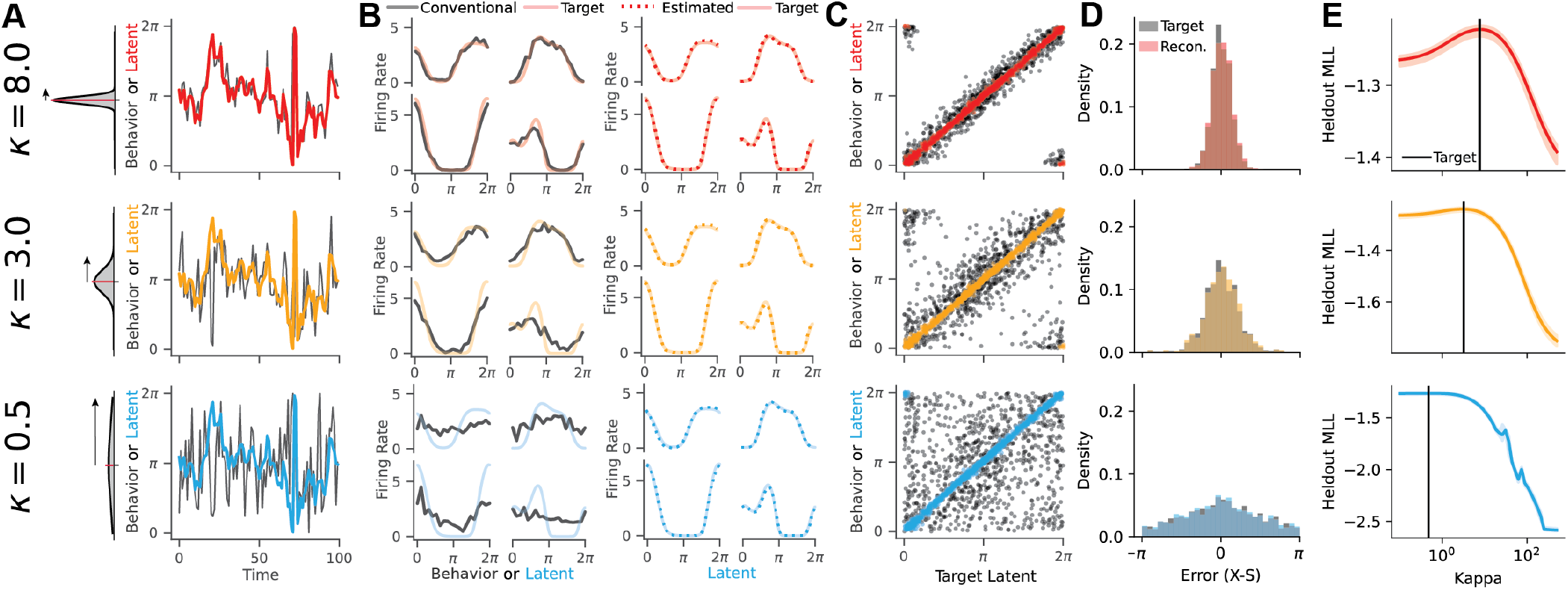
Validation on Simulated Data. A) Model applied to simulated data, N = 30, t = 1000. Data simulated at three levels of representational fidelity: *κ* = 8.0 (Top), *κ* = 3.0 (middle), and *κ* = 0.5 (bottom). Black lines indicate observed behavior and colored lines indicate ground truth latent, which is identical across simulations. B) (left) Ground truth tuning to the latent (shaded line) plotted against conventional tuning curves (black line), and (right) against model estimates of tuning curves (dotted line). C) Scatter plot of reconstructed latent against ground truth latent (colored dots) or observed behavior (black dots). D) Distribution of representation errors (**x**_**true**_ − **s**) compared to the errors between the reconstructed latent and observed behavior 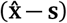. E) The marginal log-likelihood of held-out data peaks at the generative value of *κ* (black line).

When fit to these simulated datasets, the EIV model is always able to recover a precise estimate of the latent tuning curves (fig. 2B, colored lines) as well as the latent representation at each time point (fig. 2C, colored scatter plots), even though the relationship between ***s*** _*t*_ and ***x*** _*t*_ is unreliable (fig. 2C, black scatter plots). Because ***s*** _*t*_ is experimentally measured and the EIV model accurately predicts ***x*** _*t*_, we are also able to estimate the representational mismatch, ***ϵ***_*t*_ = ***x*** _*t*_− ***s*** _*t*_, on a moment-by-moment basis (fig. 2D). We will later leverage this in experimental recordings to model how latent representational errors evolve over time.

The EIV framework effectively interpolates between supervised and unsupervised latent variable models via the hyperparameter *κ*. In high fidelity regimes, the EIV model behaves like conventional tuning curve analysis, since, as *κ* → ∞, we have ***x*** _*t*_ = ***s*** _*t*_ under the model. Conversely, in low fidelity regimes the EIV model assumes that ***x*** _*t*_ and ***s*** _*t*_ are independent, since, as *κ* →0, the conditional distribution of ***x*** _*t*_ given ***s*** _*t*_ is uniform over all orientations. In this case, the model identifies a nonlinear manifold directly from the neural data in an unsupervised manner behaving like a Gaussian Process Latent Variable Model (GPLVM; Lawrence 2005, see *Methods*). For intermediate fidelity regimes, the EIV model will appropriately weight information from ***s*** _*t*_ in order to infer ***x*** _*t*_ .

In Figure 2E, we demonstrate that the fidelity parameter, *κ*, is readily identifiable through cross-validation. Across all simulated regimes, we find that the value of *κ* which maximizes the held out log likelihood closely matches the ground truth. This provides a data-driven strategy for tuning the EIV model on experimental data, where the degree of representational fidelity is unknown *a priori*.

### 2.3 Application to Head Direction Coding

Neural encoding of position and head direction is impaired when sensory cues are removed (Zhang et al. 2014; Chen et al. 2016). To investigate the ability of EIV regression to quantify these impairments, we first analyzed recordings from mouse anterodorsal thalamic nucleus (ADn), a region with a robust head direction signal (Taube 1995; Peyrache et al. 2015). In each experimental session, we analyzed three contiguous epochs (fig. 3A). In the first epoch, the mouse navigated freely in a well-lit environment with full sensory information. In the second epoch, it navigated the same space in darkness, lacking visual cues. In the final epoch, the mouse was head-fixed and placed on an air-floating track that rotates beneath it, disrupting vestibular information while preserving visual cues about orientation (Go et al. 2021; Stuart et al. 2024; Carrillo Segura et al. 2026). The full dataset consisted of 13 experimental sessions recorded across 5 mice, with the number of neurons per session ranging from *N* = 18 to *N* = 256.

**Figure 3.**
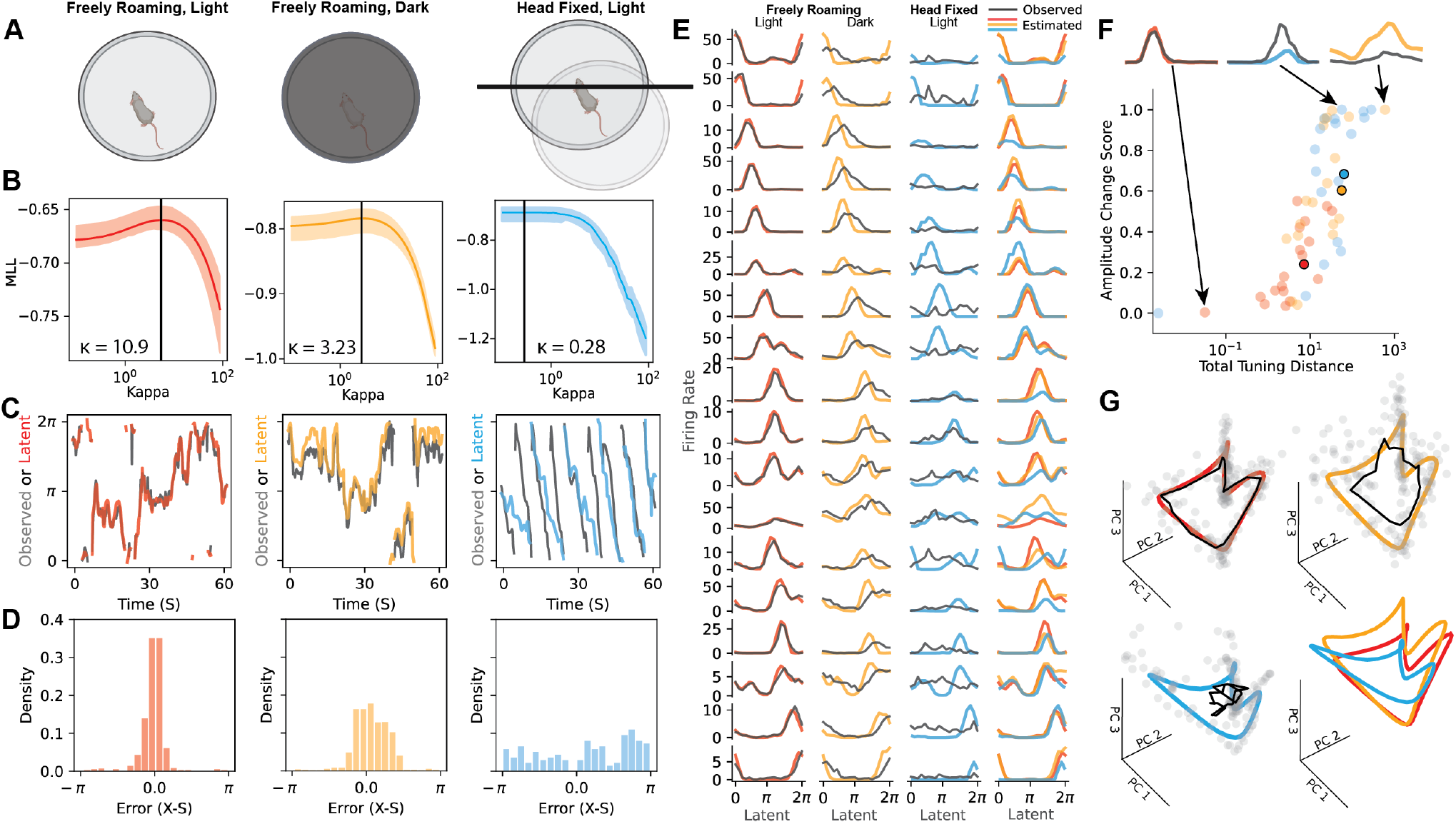
EIV Model Recovers Tuning in Mouse ADn Across Perceptually Limited Behavioral Conditions. A) Task schematic - in each session, the mouse navigates a circular environment in three distinct perceptual settings: freely roaming & light (left), freely roaming & dark (center), head fixed & light (right). B) Model performance across a range of values of *κ* for each behavioral epoch, reported as the marginal log likelihood of held out data over 10 random folds of testing and training sets. Black line indicates best performing value of *κ*. C) Observed (black) and latent (color) head direction reconstructed from model estimated tuning-curves. D) Distribution of representational mismatch *ϵ*_*t*_ over each epoch. Note the concentration of *ϵ*_*t*_ around 0 for the light and dark settings, while the distribution in the head-fixed setting appears uniform. E) Empirical tuning curves estimated using observed head direction (black) and model estimated tuning curves (color) for all 18 neurons in this session. Model results become rotationally invariant as *κ*→ 0 (unsupervised regime). For the purposes of this visualization, the tuning curves for the full population are rotated uniformly to align with the observed tuning curves in the freely roaming light epoch. All three epochs are plotted against each other in the far right column. F) Changes in the EIV tuning curves (*f*) across epochs are characterized relative to the conventional tuning in the well-lit setting (*g*) using two metrics - total distance (|| *f* − *g*||) and gain distance 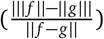. These changes are qualitatively demonstrated by example tuning curves with nearly no difference from freely roaming light (red), mostly gain change (blue), and both profile and gain change (yellow). Epoch mean across neurons shown with black outline. G) Full set of inferred tuning curves projected into same PCA space (color), alongside observed tuning curves (black) and a sample of timepoints projected into that space (gray dots). Refer to conceptual demonstration in (Figure 1 G-I).

For each session, we separately fit EIV regressions to the well-lit, dark, and head-fixed behavioral epochs. The EIV model was constructed as in fig. 2, where ***s*** _*t*_ represents the noisy measured heading direction and ***x*** _*t*_ represents the internal representation of heading direction. As before, the representational fidelity is quantified by a parameter, *κ*, which was tuned by cross-validation.

Figure 3 shows results on a representative session. In this session, we observed that the representational fidelity identified by cross-validation decreases across the well-lit, dark, and head-fixed epochs (fig. 3B). This was associated with progressively increasing deviations between the inferred latent representation and externally measured head direction (fig. 3C-D). These results confirmed our expectation that mismatch between internal representations and external measurements increase as sensory cues are removed or disrupted. Furthermore, head fixation led to a stronger effect than turning off the lights. In the head-fixed session, the mouse favors one direction of motion, while the latent representation moves in the same direction but at a different speed than the observed behavior (fig. 3C, right, blue versus black lines).

As expected, conventional tuning curves showed strong tuning for head direction in the well-lit condition, but flatten out in dark and head-fixed conditions (fig. 3E, black lines). In contrast, EIV-derived tuning curves were, in many cases, stable across the three conditions (fig. 3E, colored lines). Some neurons are exceptions to this trend, showing marked differences in tuning under head fixation. To quantify these differences, we compared the EIV tuning curves (*f*) with tuning curves estimated from the well-lit epoch (*g*) using two geometric measures: *total distance* (|| *f* − *g*||), which captures overall changes in the response profile, and *gain distance* 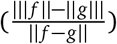, which captures changes in the matched response profile accounted for by gain modulation. Higher values on the x-axis indicate more change in the tuning curve. When the values fall near the top of the y-axis, this change can largely be attributed to a change in tuning curve amplitude rather than a more complex change in tuning profile (fig. 3F).

These findings support a view in which the primary effect of sensory deprivation is a decoupling of the latent manifold from external covariates, rather than a degradation of the manifold itself. Neural tuning to the latent representation, ***x*** _*t*_, is mostly preserved, meaning that neural activity lies close to a similar 1D ring manifold across all three experimental conditions. At the level of individual neurons, however, we observe some gain modulation that is heterogeneous across the population, and would be obscured by traditional regression based approaches.

Goodness-of-fit criteria for the EIV model are also comparable across the three conditions (pseudo-*R*^2^: light = 0.686 ± 0.004, dark = 0.614 ± 0.012, head-fixed = 0.602 ± 0.017), suggesting that deviations that are locally orthogonal to this manifold are comparable across conditions. However, deviations *along* the manifold, which correspond to mismatch between the latent representation ***x*** _*t*_ and measured head direction ***s*** _*t*_, differ considerably across the three conditions—they are small under well-lit conditions, modestly large in the dark, and extremely large under head fixation. These trends are visible in PCA projections of the raw data and manifolds derived from the EIV model (fig. 3G).

### 2.4 Dynamics of Head Direction Residuals and Variability Across Sessions

In the last section, we used EIV regression to characterize ADn coding for head direction in a single experimental session encompassing three experimental conditions (well-lit, dark, and head-fixed). We now use the EIV model to summarize the full dataset, consisting of 13 experimental sessions recorded across 5 mice. Two experimental epochs were excluded from a subset of analyses due to insufficient data under head-fixed conditions.

We began by repeating the analysis shown in fig. 3B, where cross-validation is used to select a value for *κ* in each experimental epoch. We observed significant differences in these values of *κ* across light, dark, and head-fixed conditions (fig. 4A, Friedman test, n=11 sessions, statistic=18.56, p<1e-4). In 10 out of 13 sessions, the cross-validated *κ* value was smaller in the dark condition, relative to the light condition (sign test, n=13 sessions, statistic=0.77, p=0.09). In all 11 sessions with sufficient data in the head-fixed condition, the cross-validated *κ* value was smaller in the head-fixed condition, relative to either light or dark conditions (sign test, n=11 sessions, statistic=1.0, p=0.001). Thus, many sessions followed the trends shown in fig. 3, in which the ADn representation and measured head direction were progressively mismatched across light, dark, and head-fixed conditions.

**Figure 4.**
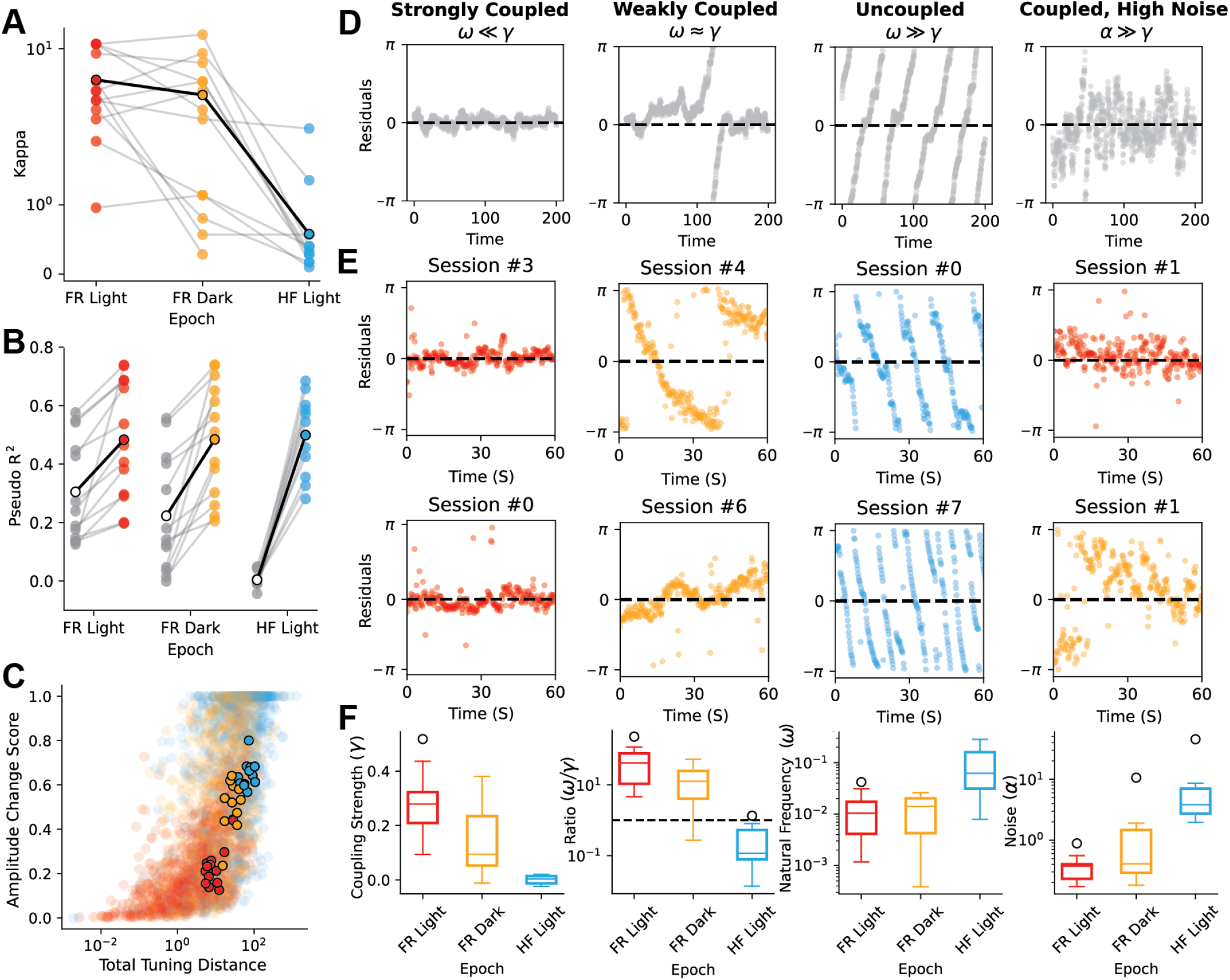
Error Dynamics of Head Direction Representations and Variability Across Animals. A) Cross-validated *κ* for each session and epoch. Black outlined points indicate mean over sessions. B) Model performance relative to a fully supervised baseline. Note that as the traditional tuning curves become more blurred (head fixed setting), the gain relative to the flat baseline nears 0 (HF Light, gray). C) Total distance in tuning from light tuning curves versus amplitude gain score for all neurons. A gain score of 1 indicates that all change can be accounted for by change in gain. Points outlined in black indicate mean for that subject/epoch. D) Simulations demonstrating four oscillatory regimes identified in the experimental data - strongly coupled, weakly coupled, uncoupled, and high noise. E) Residuals for example sessions for one minute of recording, columns indicate oscillating regimes identified in D. A residual of 0 (dotted line) indicates latent perfectly corresponds with observed behavior. F) Oscillator model parameters across sessions.

Although the fidelity of heading direction was highly variable across conditions, EIV model fits accounted for neural activity comparably well in all cases. Specifically, we quantified McFadden pseudo-R^2^ values of predicted neural firings rates using conventional tuning curves (fig. 4B, grey dots) and EIV regression models (fig. 4B, colored dots). This statistic was computed relative to a homogeneous Poisson baseline so, intuitively, a pseudo-R^2^ of zero indicates no improvement over a flat tuning curve (see *Methods*). EIV regressions predicted neural activity better than conventional tuning across all three experimental conditions, with the most pronounced improvement in the head-fixed condition where head direction coding is most error-prone. A repeated measures, two-way ANOVA applied on Aligned Rank Transformed pseudo-R^2^ values indicated significant main effects for model type (conventional tuning vs. EIV; statistic=124.90, p<1e-4) and experimental condition (light vs. dark vs. head-fixed; statistic=6.99, p=0.002) as well as a significant interaction between these factors (statistic=11.09, p=1e-4). This is consistent with the results illustrated in fig. 3F, which indicate that neural activity lies similarly close to a ring manifold across all three conditions, even though the activity along this manifold is not always registered reliably to heading direction.

While these trends are robust, we also see a remarkable degree of variability across sessions. In particular, although the cross-validated value of the fidelity parameter *κ* tends to decrease across well-lit, dark, and head-fixed conditions, we see substantial overlap across these three conditions indicating experimental condition is not the sole predictor of appropriate level of supervision. This variability in the *κ* parameter is not accounted for by differences in the number of neurons or time points analyzed (fig. S1).

EIV models readily enable us to investigate the structure of this variability. We first visualized how the model inferred residuals, ***ϵ***_*t*_ = ***x*** _*t*_ − ***s*** _*t*_, evolve over time in each experiment. Intuitively, when the ADn representation precisely tracks the measured heading direction, we would expect the residuals to revert towards zero. That is, if ***ϵ***_*t*_ > 0, then we would expect Δ***ϵ***_*t*_ = ***ϵ***_*t*+1_ − ***ϵ***_*t*_ to be, on average, less than zero. Vice versa, if ***ϵ***_*t*_ < 0, then we would expect Δ***ϵ***_*t*_ > 0, on average; deviations in internal representation are “pulled” back to the behavioral head direction by environmental signals. Alternatively, when environmental cues are limited and the ADn representation becomes decoupled from the observed head direction, the residual dynamics may be dominated by internal signals, allowing Δ***ϵ***_*t*_ to vary unconstrained by the environment. In this setting, the residuals of these two periodic signals may exhibit more autonomous oscillatory behavior.

To capture the dynamics of this error signal over time, and characterize switching between these coupled and decoupled regimes, we fit a simple oscillator model to the residuals (see *Methods*). This oscillator model is comprised of three parameters: the natural frequency ***ω***, which reflects the intrinsic oscillatory frequency of the residuals, the coupling term ***γ***, which captures the strength of the pull of deviations back to 0, and an additive noise term ***α*** which can drive transitions between these dynamical states. Simulated examples demonstrating interactions between these parameters can be found in Figure fig. 4D, alongside representative examples from the experimental data (fig. 4E).

The distribution of these fitted parameters over sessions is summarized in (fig. 4F). When the coupling term ***γ*** far exceeds the natural frequency ***ω***, the latent and observed behavior remain locked (fig. 4D-E, left). Across sessions, the coupling strength ***γ*** decreases in the dark environment and is weakest in the head fixed setting (fig. 4F, left), while natural frequency remains near 0 in the light and dark settings and increases substantially in the head fixed condition (fig. 4F, third column). When ***γ*** ≈ ***ω, s*** _*t*_ and ***x*** _*t*_ exhibit weak coupling, with intermittent oscillations or sustained deviations (fig. 4D-E, second column). The ratio 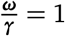 represents the critical transition point between coupled and decoupled states, (fig. 4F, second column), with nearly all head-fixed epochs falling below this threshold and most light and dark sessions exceeding it. Although noise levels increase in the dark and head-fixed epochs, sessions can maintain coupling with sufficiently large ***γ*** (fig. 4D-F, fourth column).

### 2.5 Application to Grid Cell Representations

We additionally applied EIV regression to recordings of grid cell ensembles in rat MEC released by Waaga et al. (2022). The dataset consisted of five sessions collected from four animals, with *N* = 83-258 cells per session across 2-3 grid modules. In these experiments, rats navigate a circular open-field environment first in the dark, then in well-lit conditions with an orienting cue. Although there is no head-fixed condition, the structure of this experiment otherwise mirrors the ADn dataset investigated in Figures 3 and 4, motivating the use of EIV regression. Importantly, the positional tuning curves in this dataset are multimodal over 2-dimensional space, thereby providing a more difficult test case for our methodology. Since position is not a periodic variable, we truncate the range of the stimulus and latent space to enable the EIV model, as previously described, to learn non-periodic tuning functions (see *Methods*).

As in earlier analyses, we fit the light and dark epochs separately. Across both conditions, we find that the EIV model recovers sharpened tuning curves relative to conventional rate maps that are blurred or even nearly uniform (fig. 5A-B). Similar to the head direction setting, the optimal value of *κ* for each epoch and session indicates that the error in the latent representation is significantly higher for recordings from the dark environment, relative to the well-lit condition (fig. 5C). Across all sessions, the standard deviation of the residuals between the model-inferred latent variable, ***x*** _*t*_, and the measured position, ***s*** _*t*_, ranged from 13.9 - 28.5 cm and 20.15 - 32.7 cm in light and dark conditions, respectively.

**Figure 5.**
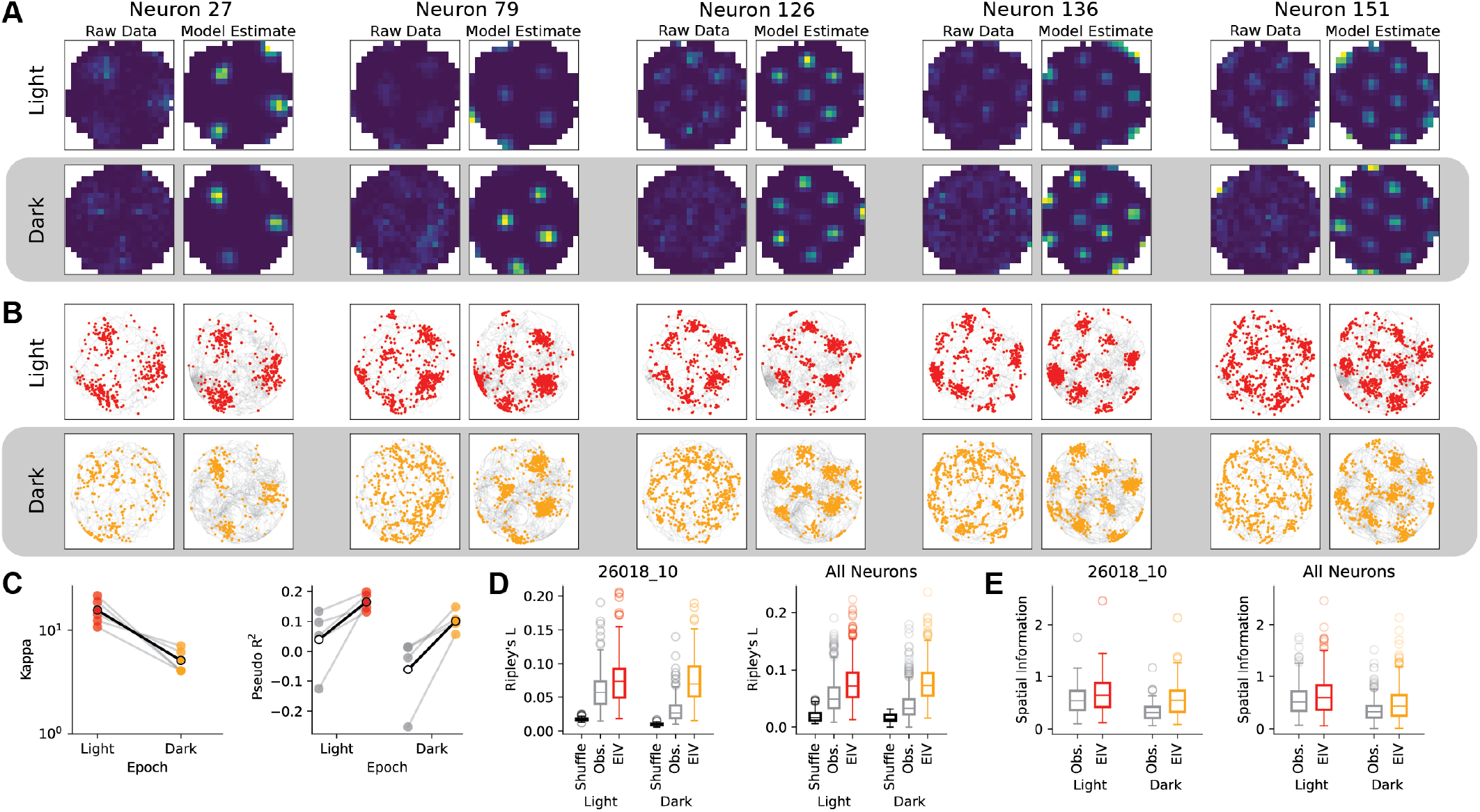
EIV Model Inference of Spatial Tuning Curves in Rat MEC Grid Cells. A) Observed (left) and EIV estimated tuning curves (right) for five example neurons from the light and dark epochs. B) 30 seconds of spiking activity from example neurons plotted relative to observed location **s**_**t**_ (left) and EIV latents **x**_**t**_ (right). C) Optimal values of supervision for light and dark epochs for each of the five rat, as well as the improvement in pseudo-*r*^2^ for the EIV model (colors) relative to a homogeneous Poisson baseline (gray). D) Spatial information for example rat (left) and for all neurons pooled over sessions (right). E) Ripley’s L for the example rat shown in A and B (left, rat #26820), as well as for all neurons pooled over sessions (right).

To further quantify these effects, we considered two metrics: spatial information and Ripley’s L. Spatial information is a widely used measure of the quantity of information a neuron’s firing activity conveys about the animal’s position in the environment (Skaggs et al. 1996). This metric recovers greater information content from the EIV tuning curves than the observed rate maps (fig. 5D), suggesting that neural activity carries more information about the model-inferred latent variable than measured position. Spatial information is sensitive to choices of bin size, and can be artificially inflated due to low firing rates (Treves and Panzeri 1995). To supplement this metric, we used Ripley’s L as a measure of spatial clustering, quantifying deviation from random distribution at increasing spatial scales, with larger values indicating greater clustering (Møller and Waagepetersen 2017). Figure 5E shows Ripley’s L both for the example rat from (fig. 5A, B) and for all neurons across subjects. Spatial clustering is lower in the dark epochs when estimated using spikes relative to the observed behavior **s**_**t**_ for the dark epochs, but comparable across light and dark epochs when evaluated using the spikes relative to the EIV-estimated latent **x**_**t**_.

We further compared the performance of EIV models with the optimal level of supervision (*κ* tuned by cross-validation) to models with full supervision (i.e. *κ* = 1000) and lacking supervision entirely (i.e. *κ* = 0.1) in an example well-lit session. Following the procedure in (George et al. 2024), we initialize the tuning curves of the unsupervised model with the supervised tuning curves so that the latent space bears some resemblance to the animal behavior. As before, fully supervised models yield spiking activity that is spatially dispersed (fig. 6A, top), while the EIV model yields tight spatial clusters of spikes (fig. 6A, middle). Interestingly, while fully unsupervised models also yield clear spatial clusters of spikes, these spike patterns diverge considerably from the original behavior. In particular, since these models have a preference for smooth tuning curves under the prior, they tend to merge spatial firing fields together and result in a decreased number of fields (fig. 6A, bottom). Together these results illustrate clear and distinct drawbacks to conventional tuning curve analysis and fully unsupervised latent variable models, and demonstrate how EIV models can achieve a “sweet spot” in between these two extremes.

**Figure 6.**
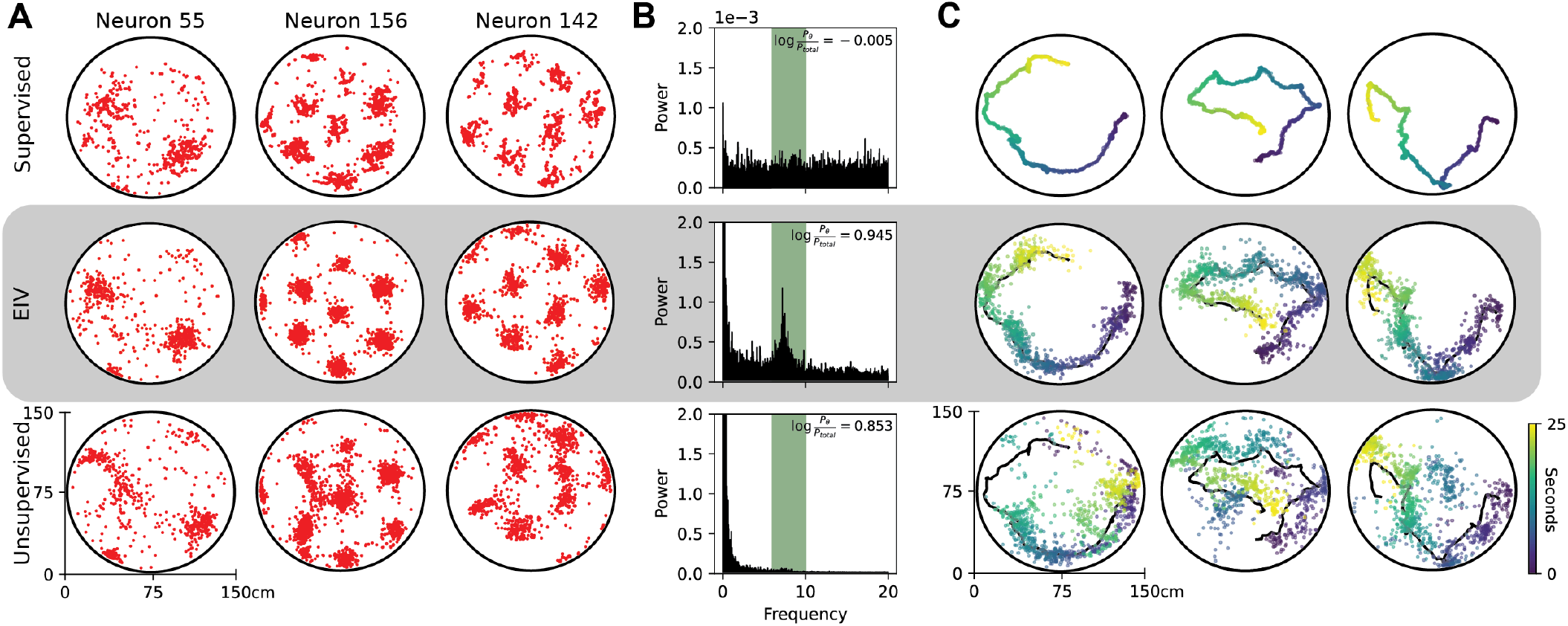
Latent Representations Inferred by Supervised, EIV, and Unsupervised Models. A) Comparison of spike time localization across three model settings - supervised (*κ* = 500), EIV (optimal *κ*), and unsupervised (*κ* = 0.1) in an example light epoch. B) Normalized power spectrum of model residuals (**x**_**t**_ − **s**_**t**_) across the three model settings. Theta frequencies (6 − 10*Hz*) are highlighted in green. C) Position observations (black) and model-inferred latent representations (colored points) over three 25 second windows across the three model settings - color indicates timepoint in window.

We next investigated the residuals of the model-inferred internal representation, given by the latent ***x*** − _*t*_, and the observed behavior, given by 2D position ***s*** _*t*_ . By construction, these residuals are uninteresting in the supervised model (fig. 6B, top), which enforces ***x*** _*t*_ − ***s*** _*t*_ ≈ 0. In the EIV model and fully unsupervised model, these residuals capture the magnitude of mismatch between the grid cell representation of position and the experimentally measured position at each moment in time. In both cases, we find that the magnitude of the residuals fluctuate in the theta rhythm frequency band (6-12Hz), consistent with a variety of previous observations in navigational circuits (Lisman and Redish 2009; Vollan et al. 2025). However, the modulation of the theta rhythm is much cleaner and distinguishable in the EIV model (fig. 6B, middle), compared with the fully unsupervised model (fig. 6B, bottom) (log ratio of theta power to baseline for Supervised 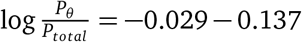, EIV 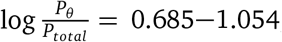, Unsupervised 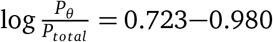). We interpret this as further evidence that the unsupervised model’s latent space diverges substantially from the measured behavior, harming interpretability.

These trends are visible when plotting 2D trajectories of the animal’s position against model-inferred latent representations. The supervised model inflexibly enforces the latent representation to equate with the measured position at all time points (fig. 6C, top). The unsupervised model is overly flexible, allowing the latent to essentially decouple from the measured behavior (on average, the residual in the light (dark) is 31.6 cm (26.6 cm), representing 19.2% (28.3%) of the arena). The EIV model with *κ* tuned by cross-validation provides an intermediate level of flexibility, allowing moderate differences between position and the latent representation (20.7 cm (33.9 cm), on average, representing 12.5% (20.6%) of the arena).

### 2.6 EIV position decoders are more accurate and consistent

As a final analysis, we examined how EIV models can be leveraged to decode neural representations on held out data, even in the complete absence of behavior. To do this, we revisited classic Bayesian decoding methods, which model the density of the represented variable, ***s*** _*t*_, as proportional to the likelihood of neural activity given ***s*** _*t*_ times a prior density. In the context of animal navigation, this scheme has been used to infer representations of position during sleep (Louie and Wilson 2001; Carr et al. 2012) and periods of immobility (Foster and Wilson 2006; Dragoi and Tonegawa 2011; Karlsson and Frank 2009; Pfeiffer and Foster 2013), which have been interpreted as neural replay or preplay of experiences.

Somewhat curiously, classical Bayesian decoders are trained under the assumption that the neural circuit faithfully represents ***s*** _*t*_ but then, at test time, reverse this assumption completely. Put differently, classical Bayesian decoders are constructed from supervised tuning curves, but at evaluation time they allow the decoded position deviate from ongoing behavior (e.g. to find remote replay events). EIV models make these assumptions explicit and consistent across model training and evaluation by introducing a latent representation, ***x*** _*t*_, that is similar (but not necessarily equal) to ***s*** _*t*_ . After fitting the model, we can predict the posterior density of ***s*** _*t*_ by marginalizing over ***x*** _*t*_ given held out neural observations. Alternatively, we can treat the posterior density of ***x*** _*t*_ given neural observations as indicating the decoded position. In either case, it is easy to show that we recover conventional Bayesian decoders in the limit that *κ* → ∞ (see *Methods*).

We reasoned that, by adaptively tuning *κ* to each dataset, EIV models could outperform conventional Bayesian decoders. We have seen that if *κ* is chosen too large, tuning can be obscured (see e.g. fig. 6A, top vs. middle sub-panels). On the other hand, if *κ* is too small, then the inferred latent variable becomes decoupled from the behavior (see fig. 6C, middle vs. bottom sub-panels). Thus, one might expect that intermediate values of *κ* may provide the best decoding performance.

We examined this prediction in the dark epochs of the grid cell dataset by evaluating the ability of EIV models to decode position on held out data as a function of *κ* (fig. 7A). We found that decoding accuracy peaked at intermediate values of *κ* on each of the five animal subjects, suggesting that a well-tuned EIV model can often outperform classical Bayesian decoder models (fig. 7B). In the well-lit environment, where the observed behavior serves as a more reliable proxy for the latent, the EIV tuning curves offer only a modest change relative to traditional Bayesian decoding (fig. S2D) Importantly, the animal’s position at ***s*** _*t*_ is never provided to any of the models at test time.

**Figure 7.**
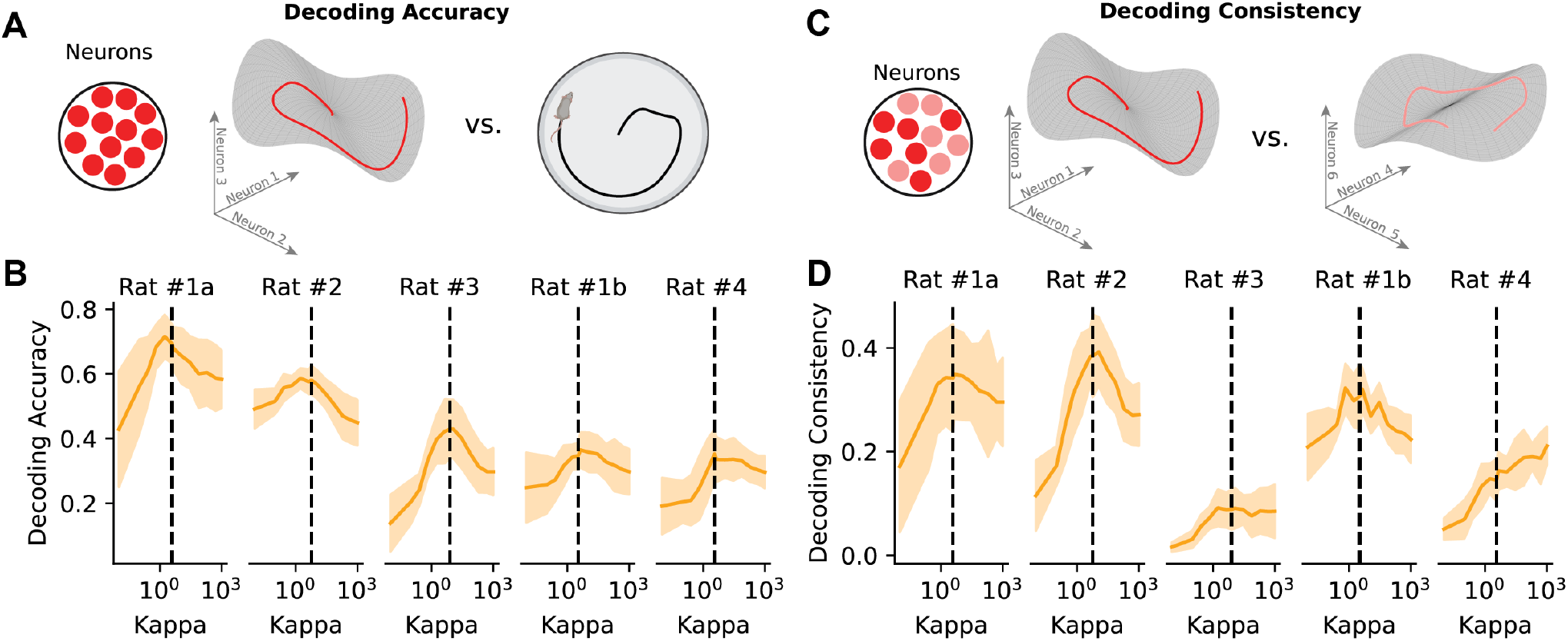
Connections between EIV Regression and Bayesian Decoding. A) Schematic of decoding accuracy analysis. Decoding accuracy compares latents reconstructed from the EIV-estimated tuning curves for the full population to the observed behavior. B) *R*^2^ decoding score indicating alignment between latent estimates and observed behavior for EIV estimated tuning curves, dashed line indicates optimal *κ*. C) Schematic of consistency analysis. Decoding consistency compares latents reconstructed from the EIV-estimated tuning curves of two equally sized, randomly selected subpopulations from the full set of recorded neurons. D) *R*^2^ latent reconstructions from each subpopulation across range of *κ*, dashed line indicates optimal *κ*.

The above result assumes that the goal of decoding is to infer the animal’s true position, ***s*** _*t*_, during behavior. However, when the scientific goal is to analyze preplay or replay of experiences, a decrease in decoding accuracy is not necessarily indicative of a failure in model performance, but instead meaningful mismatch between behavior and representation. We therefore performed a complementary analysis in which we randomly partitioned co-recorded neurons into two sub-populations and evaluated whether model-inferred latent trajectories were consistent across the two sub-populations (fig. 7C). We reasoned that a model could be highly consistent by this metric—and therefore amenable for scientific interpretation—without necessarily having strong decoding performance. Under this metric, intermediate values of *κ* still performed best in three out of five sessions (fig. 7D). In the remaining two sessions, intermediate values of *κ* performed comparably to conventional Bayesian decoding (*κ* → ∞) and outperformed unsupervised models (*κ* → 0).

## 3 Discussion

We introduced a general-purpose method based on EIV regression to characterize the degree of mismatch between neural representations and external sensory or behavioral measurements. Our work bridges a fundamental gap between the two dominant paradigms in systems neuroscience: supervised tuning curve analysis and unsupervised latent variable models (LVMs), both of which possess distinct limitations.

In particular, conventional tuning curves ignore the reality that sensory inputs are often obscured or noisy, and that internal representations may decouple from the environment during cognitive processes like planning or navigation. Even under favorable conditions (e.g. navigation in well-lit environments), internal neural representations may imperfectly track external variables, resulting in tuning curves that are over-smoothed or “smeared out” (see 5A-B, top rows). The magnitude of this effect likely depends on experimental conditions (e.g. recording in the light vs. dark) and behavioral state (e.g. whether the animal is highly engaged or disengaged from the behavioral task).

Conversely, while unsupervised LVMs are powerful for capturing structure in high-dimensional neural spike trains, they neglect the rich experimental context available to the researcher. Often, unsupervised neural latents are inferred only to relate them back to behavioral covariates through ad hoc analyses. This two-step approach is not only sub-optimal from a statistical perspective, it is often under-constrained. Indeed, even when latent variables are initialized in a supervised manner, they often diverge from their intended behavioral correlate after the model is fit (see Fig. 6).

EIV regression can be viewed as a partially supervised method where a continuous hyperparameter, *κ*, tunes the degree of supervision and represents the fidelity of the neural representation. As *κ*→∞, the internal representation perfectly tracks the external variable of interest and EIV regression converges to a GLM tuning curve model with nonlinear basis functions. Conversely, as *κ* → 0, the internal representation is completely decoupled from externally measured variables and EIV regression converges to an unsupervised manifold learning method called a Gaussian Process Latent Variable Model (GPLVM; Lawrence 2005). Importantly, *κ* can be tuned in a data-driven fashion via cross-validation, enabling it to be used as a statistic for downstream analysis. For example, in ADn and MEC populations, we not only observed that *κ* decreased when sensory information was disrupted (as expected), we additionally observed substantial animal-to-animal variability in *κ* that was not easily explained by confounding differences (e.g. size of the recorded neural population) (fig. S1). These individual differences are difficult to quantify without a statistical model, such as EIV regression, that is specifically tailored to address them.

Our method extends a growing body of work that incorporates behavioral labels into latent variable models of population activity. Several recent methods extend traditional approaches to allow for refining model trajectories beyond an initial model fit (George et al. 2024), explicitly incorporating correlations between neurons into GLMs (Vollan et al. 2025; G. Wei et al. 2024), or leveraging behavioral labels during model fitting (Schneider et al. 2023; Zhou and X.-X. Wei 2020). While these are powerful methods, they are often ambiguous about the degree to which behavioral labels determine the structure of the latent space. In EIV regression, this dependence is explicit and can be inferred in a statistically principled manner.

In addition to quantifying overall representational fidelity through cross-validated estimates of *κ*, EIV regression provides moment-by-moment estimates of the latent representation and its residual with respect to external measurements. This enabled us to build a simple dynamical model which captured the transient decoupling and recoupling of ADn representations of head direction to external measurements of heading (see Fig. 4). This revealed a variety of dynamical regimes across experimental conditions and subjects, including (a) high-precision, high-accuracy representations that tightly tracked the true head direction, (b) high-precision, low-accuracy representations that intermittently drift away from the measured head angle, and (c) low-precision, high-accuracy representations that are noisy from moment-to-moment but are largely centered at the measured head angle. These results provide rich, quantitative, and novel constraints for attractor-based circuit models of the mammalian head direction system. In principle, this approach could be applied to study the continuous dynamics of a variety of other sensory or motor errors beyond navigation, such as working memory (Wimmer et al. 2014) and visual tracking tasks (Bonnen et al. 2015).

Another useful feature of EIV regression is its ability to disentangle changes in neural tuning from experimental noise or perceptual mismatch. In fig. 1, we simulated a simple example where the underlying manifold of neural representations is unchanged across three conditions (see panels g-i), while noise in the latent representation causes conventional tuning curves to flatten (see panels d-f). Neuroscientists commonly report similar changes in tuning shape—e.g. in animal models of Alzheimer’s disease (Zhao et al. 2014) and in early sensory representations under metabolic constraints (Orbán et al. 2016). However, it is important to remember that these effects can be explained in at least two ways, with very distinct circuit-level implications. First, individual neurons may each become more noisy or less responsive to the signal of interest (i.e., a decrease in precision). Alternatively, as illustrated by the simulation in fig. 1, the properties of individual neurons might remain completely intact, while the alignment between the latent circuit representation and measured variable degrades (i.e., a decrease in accuracy). For the two navigation datasets analyzed in this study, our analysis strongly favors the latter explanation over the former—tuning to the inferred latent variable was remarkably stable across conditions, aside from some modest gain modulation in ADn under head fixation (fig. 3, 4).

Together, these results demonstrate that EIV regression provides a flexible and practical framework for tracking internal neural representations in relation to externally measured variables. The model yields several interpretable quantities, such as the representational fidelity parameter *κ*, tuning curves to the latent representation, and a moment-by-moment time course of residuals, all of which can be compared across conditions, tasks, and subjects. We anticipate that applying this framework to additional brain areas and behavioral paradigms will help clarify when and how internal neural estimates decouple from external variables and refine theories of how population codes support flexible cognition in noisy, partially observed environments.

## Acknowledgments

We thank Simón Carrillo-Segura, José Hurtado, Dora Angelaki, and André Fenton for sharing and discussing the ADn head direction dataset with us. We are also grateful to Guillaume Viejo for insights into the navigation system, and his contributions as part of the Flatiron Institute NeuroRSE team, alongside Edoardo Balzani, Billy Broderick and Sarah Jo Venditto, for their feedback and assistance with code development. This work was supported by a McKnight Fellowship and NIH funding sources (1RF1MH133778-01).

## Generative AI Disclosure

During the preparation of this work the authors used Claude, Perplexity, and ChatGPT to prototype small portions of code and to edit sentences of the text for clarity. After using this tool/service, the authors reviewed and edited the content as needed, and take full responsibility for the content of the published article.

## 4 Methods

### 4.1 Experimental Model and Subject Details

#### 4.1.1 Simulated Data

To validate that we can recover the true, generative value of *κ*, we explored three synthetic datasets with known values of *κ* = 0.5, *κ* = 3.0, and *κ* = 8.0. Each dataset consisted of 30 neurons and 1000 timesteps. The spiking activity (and therefore the latent) was identical across all three settings - only the behavioral observations are changing. For the purposes of visualization, the latent evolves according to a periodic random walk, although no dynamics were used in the fitting of this model. For model fitting, candidate values of *κ* were evaluated on a log spaced grid from 0.1 to 500. Tuning curves were drawn from the Gaussian Process Prior, with a length scale *l* = 0.25 and and *σ* = 100, not dissimilar from the parameters used for the head direction data later in the paper.

#### 4.1.2 Mice Head Direction

For the head direction dataset we analyzed (fig. 3-4) shared with us from (Carrillo Segura et al. 2026). Subjects were C57Bl6/J mice, 3 to 6 at the time of the experiment. All experimental procedures were carried out in accordance with US National Institutes of Health guidelines and approved by the New York University Welfare Committee (UAWC). All efforts were made to minimize the number of animals used.

#### 4.1.3 Rat Grid Cells

For the grid cell analyses (Fig. 5-7), we analyzed publicly available data published in (Waaga et al. 2022). Subjects were male Long Evans rats, 2-4 months old at the start of the experiment. All procedures were performed in accordance with the Norwegian Animal Welfare Act and the European Convention for the Protection of Vertebrate Animals used for Experimental and Other Scientific Purposes.

### 4.2 EIV Framework

Our model relates an input covariate (behavior or stimulus) using an Error-in-Variables (EIV) framework (Carroll et al. 2006). At the core of this approach is the assumption that the measured covariate *S*_*t*_ is a noisy realization of an underlying latent state *x*_*t*_ :

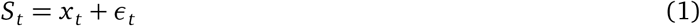

where *ϵ*_*t*_ reflects our uncertainty between the latent representation and the observed covariate. Under this framework, both the neural population activity **Y** and the behavioral measurements **S** are conditioned on the hidden latent trajectory **x**_1:*T*_ . The joint distribution over the observations, the latent variables, and the mapping weights **W** is given by:

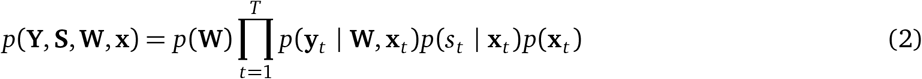

Where *p*(**W**) is the mapping prior that enforces smoothness on the tuning curves, *p*(**y**_*t*_ **W, x**_*t*_) is the neural likelihood characterizing the mapping from the latent state to observed spike counts, and *p*(*s*_*t*_ **x**_*t*_) is the behavioral likelihood accounting for uncertainty in the measured covariate. By explicitly modeling the covariate as a noisy observation, the framework accounts for errors in the behavioral signal that could otherwise bias the estimation of the neural mapping functions.

#### 4.21 Behavioral Observation Model

The specific form of *p*(*s*_*t*_ | **x**_*t*_) depends on the topology of the latent representation. For periodic variables (e.g., heading angle), we model the observation noise using a von Mises distribution:

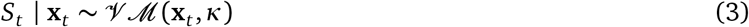

where *κ* is the concentration parameter. For non-periodic, bounded variables, the behavior and latent states are mapped to a normalized domain (0, 0.8]. By constraining the latent range to be smaller than the 1.0-period of the underlying basis set, we allow for aperiodic boundary conditions while maintaining stable function approximation at the manifold edges.

#### 4.2.2 Spiking Observation Model

The neural population activity **y**_*t*_ is modeled as a set of independent Poisson likelihoods conditioned on the latent state and the neuron-specific mapping weights **W**:

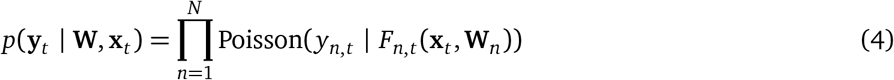

We represent the mapping from a latent manifold **x** to neural firing rates using a weight-space formulation of a Gaussian Process (GP). While GPs are typically defined by their covariance kernels, they can be equivalently expressed as a linear combination of basis functions *φ*(**x**). By taking this weight-space perspective, we avoid the *O*(*T* ^3^) complexity of matrix inversions in function-space GPs, achieving *O*(*T*) scaling with respect to the number of timepoints. This approach is well explored in the machine learning literature, where the kernel is approximated either by sampling frequencies from its spectral density (Rahimi and Recht 2007; Gundersen et al. 2021), or by using a deterministic, equispaced frequency grid (Greengard et al. 2025; Solin and Särkkä 2020), which is more closely aligned with our approach here.

Each neuron *n* ∈ {1, …, *N*} is modeled with a tuning function *f*_*n*_(**x**_*t*_) constructed from a truncated Fourier basis set. This allows the model to approximate arbitrary nonlinearities and has a formal equivalence to stationary Matérn kernels typically used in GP settings (Borovitskiy et al. 2020). The firing rate *F*_*n,t*_ is given by:

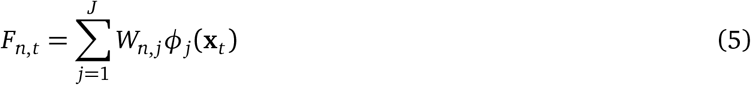

Smoothness is enforced using a frequency dependent prior on the weights of the bases, which decays at higher frequencies.

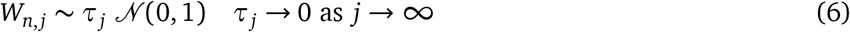

This induces smoothness by constraining higher frequency modes to have smaller weights. We threshold high frequencies at a reasonable value of *τ* = 1*e*^−3^, constraining the total number of weights we fit.

#### 4.2.3 Inference

Inference in the EIV framework requires evaluating the marginal likelihood of the observed neural activity and behavioral covariates by integrating over the hidden latent trajectory:

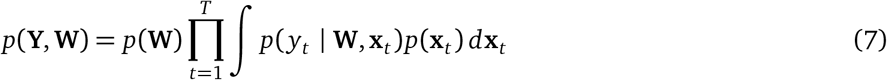

This integral is analytically intractable due to the nonlinearities introduced by the Fourier basis expansion in the spiking observation model. Therefore, we approximate this integral over the latent *x*_*t*_ using quasi Monte Carlo integration, drawing samples over the normalized input domain *x*_*t*_∈ [0, 1]^*d*^ (see Section 4.3 for further details).

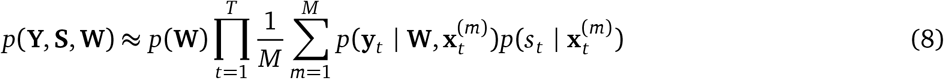

We have found that this approach was both efficient and accurate for the dimensionalities investigated here, although more complex importance sampling schemes may be needed in higher dimensional spaces. For experimental designs with non-periodic/bounded dimensions, the behavior was normalized to a spatial domain smaller than the tuning curve (0,.8], and the sampling was likewise constrained, allowing aperiodic boundary conditions.

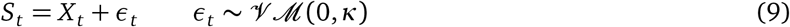

#### 4.2.4 Connections to Existing Modeling Frameworks

At its extremes, this modeling framework reduces to models familiar and broadly used in the neuroscience community. Standard PCA can be reframed as a probabilistic problem by introducing Gaussian noise in the observation model, allowing uncertainty in the latent representation and defining an explicit likelihood over the data. This perspective not only recovers standard PCA in the zero-noise limit, but also enables principled handling of noise and missing data (Tipping and Bishop 1999). Gaussian Process Latent Variable Models (GPLVMs) extend this framework by replacing the linear mapping of the latents to population activity with a nonlinear function drawn from a Gaussian process prior, which corresponds to an expansion over and infinite set of basis functions implicitly defined by the kernel *y*_*n,t*_ =∑ _*j*_ *w*_*j,n*_, φ_*j*_ (*x*_*t*_) . Under a linear kernel, the GPLVM is exactly equivalent to probabilistic PCA (Lawrence 2003). This shared probabilistic latent variable model structure forms the foundation of our project.

We extend this framework by introducing behavioral observations as noisy measurements of the latent variable. As *κ* → ∞, the noise distribution concentrates and *x*_*t*_ ≈ *s*_*t*_, recovering the fully supervised regime. In this limit, the latent is identified with the observed behavior, and with a linear kernel the model reduces to linear regression on the observed behavior *f*_*n*_(*s*_*t*_) = *w*_*n*_*s*_*t*_ . Here, inference reduces to the more straightforward, convex problem of estimating the weights **w**. More generally, nonlinear feature mappings yield generalized linear models (GLMs) of the form *y*_*n,t*_ = ∑_*j*_ *w*_*j,n*_, φ_*j*_ (*s*_*t*_).

As *κ* → 0, the noise becomes uniform, rendering *s*_*t*_ uninformative and reducing the model to a fully unsupervised latent variable model equivalent to probabilistic PCA, or its nonlinear GPLVM extension. Intermediate values of *κ* do not correspond to either extreme, but instead define a semi-supervised regime in which both neural activity and behavioral observations jointly constrain the latent representation. In this sense, our model unifies linear regression, PCA, GLMs, and GPLVMs within a single probabilistic framework.

### 4.3 Monte Carlo Samplers

Optimizing model weights requires estimating the expectation of the spiking likelihood over the latent space. These expectations are not calculable in closed form, and we therefore used two distinct Monte Carlo sampling-based approaches to take this expectation: quasi-Monte Carlo samples generated using a Roberts sequence, and sequential Monte Carlo (SMC) samples. These approaches have distinct scaling properties and assumptions about the temporal structure of the latent variables. To give some intuition on the appropriate use cases for each of these approaches, the Roberts sampler treats each time point as independent, and scales linearly with the number of temporal observations, making it efficient for larger populations of neurons and high firing regimes. In contrast, the SMC sampler explicitly models some temporal smoothness/dependence of over the latent, making it better suited to low firing regimes or small time bins, where there is less information available about the shared latent state at each timepoint. This approach introduces an additional smoothing parameter (PARAMETER), and scales worse with the window of smoothing, number of neurons and particles

#### 4.3.1 Roberts Sampler

To generate low-discrepancy samples from the latent space, we used a quasi-Monte Carlo (QMC) approach based on a Roberts sequence (Roberts 2018). Truly random MC sampling procedures, where each sample is totally independent, yields random draws which are clustered or leave gaps in regions of the sampled space. QMC mitigates this by spreading the samples more uniformly on [0, 1]^*d*^, reducing integration error and improving sampling efficiency for a fixed number of samples (Owen 2023). There are many approaches to selecting these pseudo random samples. This approach constructs a deterministic lattice on [0, 1]^*d*^, and then applies a single random offset per sample set. This yields evenly distributed samples that vary across the full latent space over iterations. We used this sampling approach to approximate the integral for the head direction and simulated datasets, using *M* = 1024 QMC samples at each timepoint to approximate the likelihood across the one-dimensional ring.

#### 4.3.2 Sequential Monte Carlo

To incorporate temporal structure in the latent trajectory, we used a sequential Monte Carlo (SMC) sampler, or particle filter, following the formulation outlined in Naesseth et al. 2019. Unlike the Roberts sampler, which treats time points independently, SMC induces some temporal dependencies by propagating Monte Carlo samples across time according to some transition model. SMC is particularly beneficial in low firing-rate regimes or when time bins are small, where individual observations provide limited information about the latent state. By maintaining dependence across time steps, the particle filter enforces smoothness in the latents and stabilizes reconstruction when instantaneous likelihoods are weak.

The algorithm is akin to sequential importance sampling, with an added resampling step. Particles are first initialized from a proposal distribution over the latent space. At each time step, particles are updated through a resample–propagate–concatenate cycle: particles are resampled according to their importance weights, propagated forward using a transition model, concatenated to the existing latent chain, and finally re-weighted to account for the mismatch between proposal and target distributions. In our implementation, the transition dynamics correspond to a Gaussian random walk, such that proposed latent states are sampled locally around the latent at the previous time point. We employ a forward filtering pass only, without backward smoothing. The computational complexity scales with the number of particles, neurons, and the smoothing window. As a result, SMC is more computationally expensive than the Roberts sampler approach, but offers improved performance for small time bins or when temporal coupling in the latent dynamics is essential.

### 4.4 Optimization

Model weights *W* are optimized according to a procedure in the family of generalized expectation maximization algorithms (Neal and Hinton 1998; Dempster et al. 1977; Caprio et al. 2025). Expectation maximization can be viewed as a block coordinate maximization of the marginal log likelihood. In this algorithm, we alternate between taking the expectation of the population firing activity over the latent space (E-step), and taking a gradient step to optimize over the model parameters (M-step). In traditional EM, both the E-step and M-step are solved exactly in closed form. Here, the M-step has no closed-form solution. While an analytical calculation of the M-step may be possible in the Gaussian case, with a Poisson noise model some iterative local optimization would still be necessary. Instead, we use here a more general form of EM, which calculates a Monte Carlo sampling based approximation of the E-step, and then given this expectation we take a gradient step in the M-direction.

Parameter updates can be performed by a number of standard gradient based optimization methods, including L-BFGS, Adam, or stochastic gradient descent, which are all implemented in the software package. All code was implemented in Python using Jax (Bradbury et al. 2018), and optimization leveraged the Jaxopt (Blondel et al. 2021) and Optax (Hessel et al. 2020) libraries. This problem is non-convex at the unsupervised (small *κ*) extreme. L-BFGS tends to converge rapidly, but is prone to convergence to local minima. Adding noise to parameter updates can mitigate this issue, as can more principled initialization strategies, including initializing weights at many different random draws from the prior, or at the weights fit by the supervised model (see George et al. 2024).

#### 4.4.1 Rotation Invariance*/*Alignment

At the unsupervised extreme, the learned manifold is not grounded to any particular observation. The grid cell model is invariant to reflection, and the head direction model is invariant to rotation and reflection. Consequently, in more unsupervised regimes the estimated tuning curves may need to be uniformly aligned to some frame of reference to enable further analysis. In order to correct for this, we aligned the representations to the observed, light conditions. For each epoch, the single rotation or reflection that maximized the correlation between the Generally, the more strongly supervised models required smaller corrections.

#### 4.4.2 Hyperparameter Selection

Hyperparameters were selected via two rounds of cross-validation. We first selected the kernel hyperparameters (lengthscale *λ* and variance *σ*), followed by the representational fidelity hyperparameter *κ*. Kernel hyperparameters were estimated in the supervised regime using data collected under optimal (light) conditions. We would like to highlight that there was some non-identifiability in the kernel hyperparameters - there are several valid combinations of lengthscale and variance that would yield comparable model performance due to trade-offs in smoothness and amplitude scaling. We also evaluated kernel parameters at an intermediate value of *κ*. This process was repeated in two representative subjects, and the parameters averaged across these subjects. The final hyperparameters were identical across subjects and epochs. For the grid cells, we fit only the highest frequency module. Distinct priors might be more appropriate for each module, but we leave this for future research.

The representational fidelity parameter *κ* was selected after fixing the kernel hyperparameters. Models were fit across a logarithmically spaced grid of values of *κ*. The bounds of this grid were defined by the values that corresponded with fully supervised and unsupervised modeling regimes. The value of *κ* that yielded the best cross-validated marginal log likelihood on held out data was selected across several random folds of the data set. This was performed on two example subjects, and the results averaged. Automated procedures or more fully Bayesian approaches for this selection process are an area of future interest.

### 4.5 Additional Analyses

#### 4.5.1 Oscillator Model

To characterize the residual errors between estimated and observed head direction, and capture switching between coupled and decoupled regimes, we fit a simple oscillator model. Let *ϵ*_*t*_ indicate the angular difference between the latent state and observation at time *t*, which is itself a periodic signal. We model the dynamics of these residuals over time as:

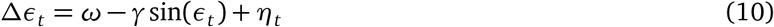

Where *ω* is the baseline drift, *γ* is the restoring pull toward alignment, and *α* indicates random fluctuations around equilibrium. These parameters were fit in two stages. First, we fit *ω* and *γ* by regressing the unwrapped derivative of the residuals Δ*ϵ*_*t*_ onto sin(*ϵ*_*t*_). We used the scikit-learn implementation of the Theil-Sen estimator to provide more robustness to outliers, particularly if the data is skewed (Pedregosa et al. 2011). Next, we estimated the noise concentration parameter on the residual angular errors *η*_*t*_ on the residuals between our fitted model and the original signal, where *η*_*t*_ ∼ *V* ℳ (0, 1*/α*). This model can be interpreted as a circular Ornstein-Uhlenbeck process, with periodic drift.

#### 4.5.2 Spatial Information

The arena was segmented into 10 cm bins. Rates and occupancy were calculated for each bin. The spatial information, in bits/second, is given by (Skaggs et al. 1996; Waaga et al. 2022):

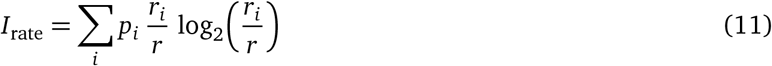

Where *r*_*i*_ is the firing rate at bin *i, p*_*i*_ the occupancy at that bin, and *r* is the average firing rate for that neuron.

#### 4.5.3 Ripley’s-L

Ripley’s-L is a variance stabilized version of Ripley’s-K statistic (Møller and Waagepetersen 2017). It operates directly on the continuous spatial coordinates of position at the time of a spike for an individual neuron, and does not require binning of the spatial signal. This statistic quantifies spatial clustering of spiking activity by estimating deviation of the number of spikes (*n*) from expected intensity of spiking (*n/A* where *A* is the area of the region) given a spatially homogeneous Poisson baseline, estimated at increasing spatial radii *r*.

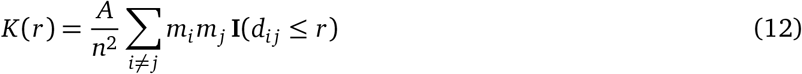

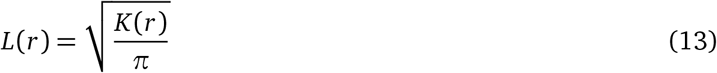

Where *m*_*i*_ and *m*_*j*_ correspond to the number of spikes at spatial locations *i* and *j*, and **I**(*d*_*i j*_ ≤ *r*) is an indicator function for points within radius *r*. Values reported are the distance from the baseline rate (*πr*^2^) at the most clustered radius.

### 4.6 Mouse Head Direction Experimental Details

For this study, we analyzed an existing dataset shared by the authors; further details can be found in Carrillo Segura et al. 2026. Mice were implanted with a 4-shank Neuropixels 2.0 probe. After experiments, probe placement in the anterior thalamus (ATN) or anterodorsal thalamic nucleus (ADN) was confirmed histologically. Recordings began three days after electrode implantation. Head-direction activity was assessed during two consecutive 8 min sessions in a circular arena (52 cm diameter): a light session followed by a dark session. The arena had of black walls with a white cue card present during the light session and removed before the dark session. To encourage exploration, food crumbles were randomly distributed prior to each session. The arena was cleaned between sessions to remove odor cues.

Head-fixed recordings were performed on the same day as freely moving sessions. Experiments were conducted using a floating arena based on the Neurotar system. The arena floats on a frictionless air cushion, allowing the head-fixed mouse to actively move the environment beneath it. This setup preserves visual, tactile, and olfactory cues during exploration while eliminating vestibular input. To encourage exploration, animals received a drop of water whenever running speeds above 5 mm/s were detected for at least 2 s. Once the mouse was head-fixed, compressed air was released to allow the arena to float. Recordings began approximately 2 min later once the platform stabilized. Animals explored the arena for 10 min. Behavioral variables were measured directly by the platform system, and these measurements provided the mouse’s X–Y position relative to the cage center and heading angle (defined as the angle between the animal’s longitudinal axis and the cage Y-axis). Across all epochs, only time points where running speed exceeded 4 cm/s were included in the analysis. Video data were analyzed offline using DeepLabCut, and extracellular recordings were spike sorted using Kilosort. Data formatting and pre-processing was performed using Pynapple (Viejo et al. 2023).

### 4.7 Rat Grid Cell Experimental Details

Rats freely explored a circular open-field arena (150 cm diameter) consisting of matte black plastic walls and a matte black rubber floor. The arena was surrounded by blackout curtains positioned 1 m from the edge to minimize external visual cues. The same arena was used for both dark and light recording sessions. For dark recordings, all light sources in the recording room were extinguished or masked. Before each session, the arena was cleaned with soap and water to minimize odor cues. Rats were introduced to the arena and allowed to forage freely for 50–60 min while small pieces of corn foam snack were scattered into the arena. Following the dark session, recordings continued for an additional 30–60 min under illuminated conditions. A single white cue card (45 cm × 150 cm) positioned on the surrounding curtains was visible from within the arena. Further experimental details can be found in Waaga et al. 2022. Data formatting and pre-processing was performed using Pynapple (Viejo et al. 2023).

### 4.8 Code Availability

The code used to generate these results is publicly available at https://github.com/neurostatslab/error-in-variables-garon-2026

## Supplementary Figures

**Figure S1.**
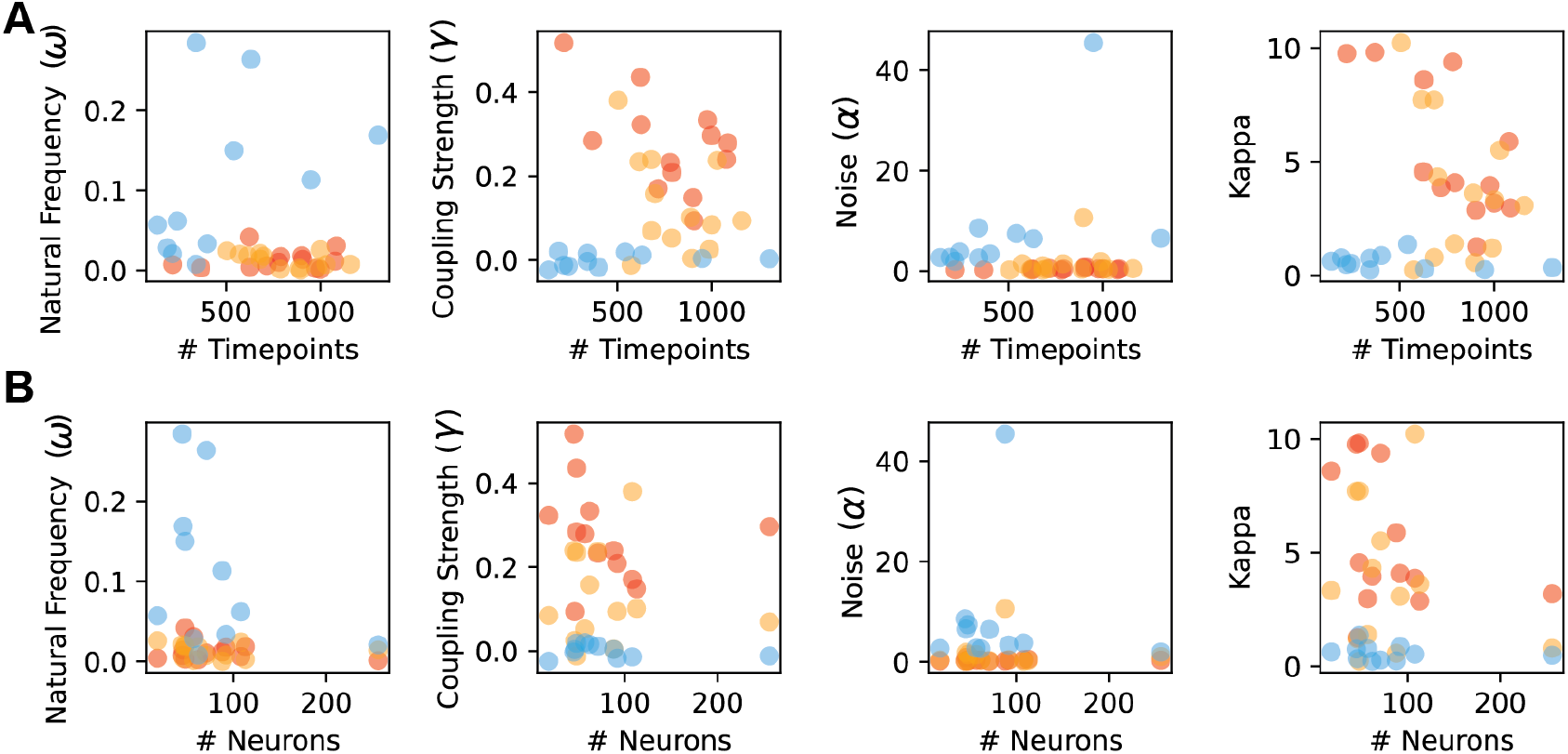
Mouse ADn - correlation between model parameters and dataset dimensions. A) Oscillator model parameters and cross-validated kappa versus number of moving timepoints for each epoch of the mouse ADn experiments. Mice tended to be stationary for longer periods of the head-fixed, light condition. B) Oscillator model parameters and cross-validated kappa versus number of neurons.

**Figure S2.**
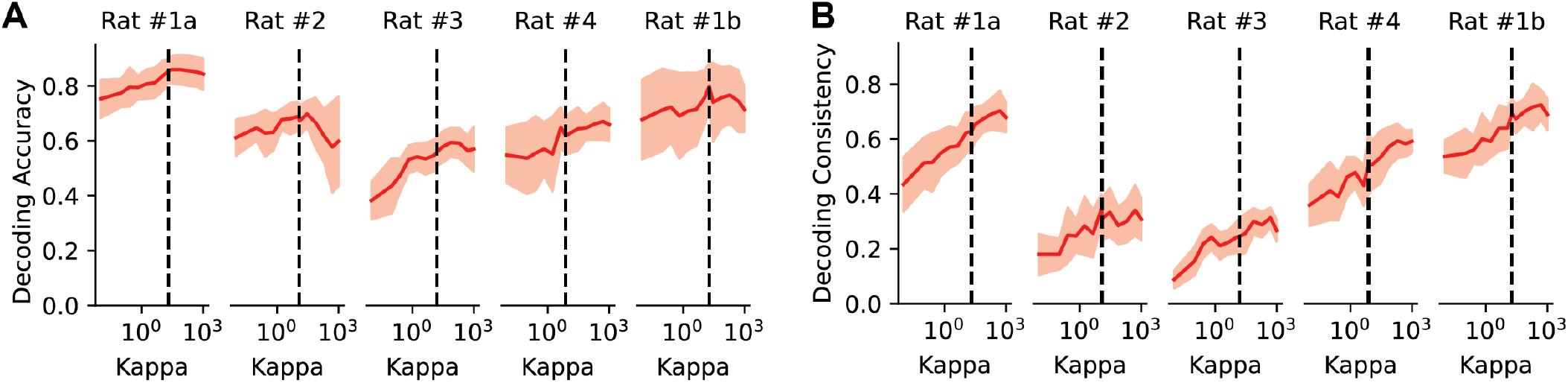
Rat MEC - Decoding accuracy and consistency for light epochs. A) *R*^2^ decoding score indicating alignment between latent estimates and observed behavior for EIV estimated tuning curves, dashed line indicates optimal *κ*. B) *R*^2^ latent reconstructions from each subpopulation across range of *κ*, dashed line indicates optimal *κ*.

## Notes

https://github.com/neurostatslab/error-in-variables-garon-2026

### Competing Interest Statement

The authors have declared no competing interest.

### Summary of Updates

Fixed several typos, updated github link.

