## Supplemental Figures for "Tracking the Fidelity of Internal Neural Representations with Error-In-Variables Regression"

### Supplementary Figures

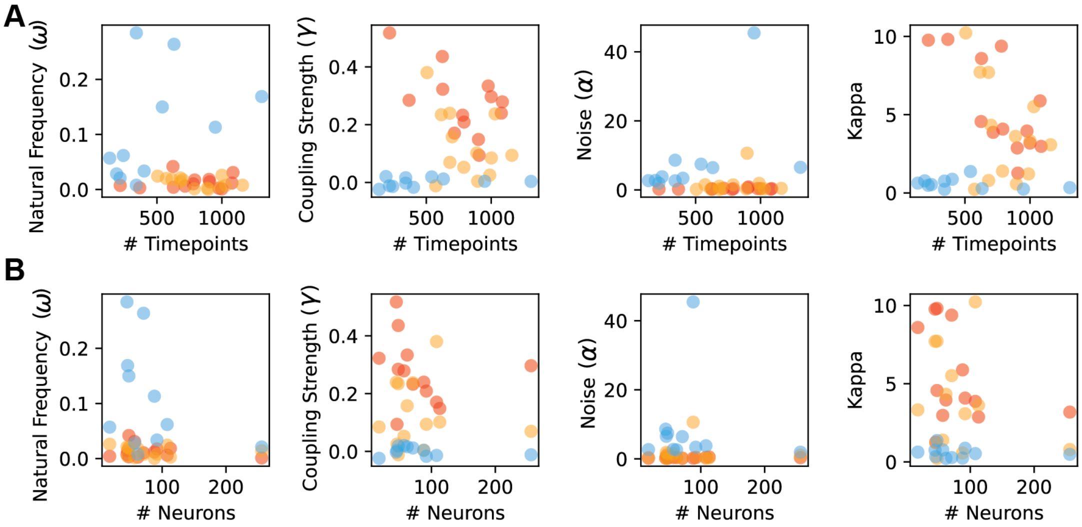


*Figure S1:* **Mouse ADn - correlation between model parameters and dataset dimensions** A) Oscillator model parameters and cross-validated kappa versus number of moving timepoints for each epoch of the mouse ADn experiments. Mice tended to be stationary for longer periods of the head-fixed, light condition. B) Oscillator model parameters and cross-validated kappa versus number of neurons.

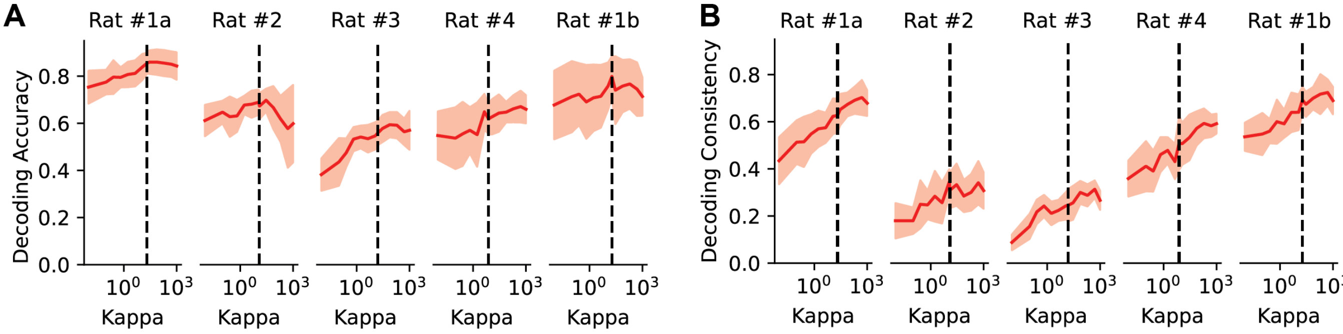


*Figure S2:* **Rat MEC - Decoding accuracy and consistency for light epochs** A) $R^2$ decoding score indicating alignment between latent estimates and observed behavior for EIV estimated tuning curves, dashed line indicates optimal $\kappa$. B) $R^2$ latent reconstructions from each subpopulation across range of $\kappa$, dashed line indicates optimal $\kappa$.
